# Direct 3D-bioprinting of hiPSC-derived cardiomyocytes to generate functional cardiac tissues

**DOI:** 10.1101/2023.06.12.544557

**Authors:** Tilman U. Esser, Annalise Anspach, Katrin A. Muenzebrock, Delf Kah, Stefan Schrüfer, Joachim Schenk, Katrin G. Heinze, Dirk W. Schubert, Ben Fabry, Felix B. Engel

## Abstract

3D-bioprinting is a promising technology to produce human tissues as drug screening tool or for organ repair. However, direct printing of living cells has proven difficult. Here, we present a method to directly 3D-bioprint human induced pluripotent stem cell (hiPSC)-derived cardiomyocytes embedded in a collagen-hyaluronic acid ink generating centimeter-sized functional ring- and ventricle-shaped cardiac tissues in an accurate and reproducible manner. The printed tissues contained hiPSC-derived cardiomyocytes with well-organized sarcomeres and exhibited spontaneous and regular contractions, which persisted for several months and were able to contract against passive resistance. Importantly, beating frequencies of the printed cardiac tissues could be modulated by pharmacological stimulation. This approach opens up new possibilities for generating complex functional cardiac tissues as models for advanced drug screening or as tissue grafts for organ repair or replacement.

## Main

One goal of biofabrication is to engineer tissues, organs, or parts of them to study or treat diseases for which there are limited therapeutic options, such as end-stage organ failure. Initially, acellular scaffolds with an architecture, topography, porosity, physicochemical properties, and mechanical properties corresponding to the target tissue were generated. However, a major disadvantage of acellular 3D scaffolds is the difficulty of colonizing them with cells, especially in an organized manner. Another approach is the casting of cell-laden hydrogels. However, this strategy does not allow the fabrication of hierarchical structures. 3D-bioprinting has the potential to solve major bottlenecks and current problems faced in tissue engineering.

Cardiovascular disease is the most common cause of death in the world^1^. In search of novel therapeutic strategies, engineered cardiac tissues are valuable tools to model disease and study drug response. In addition, pre-clinical animal studies suggest that transplantation of such engineered tissues can improve heart function after infarction^2-6^. The safety and efficacy of such a procedure for patients is currently being investigated in a clinical trial (ClinicalTrials.gov Identifier: NCT04396899). Most commonly, cardiac tissues are produced by casting a mixture of extracellular matrix (ECM) proteins and cardiomyocytes as well as other cell types of the heart. In this manner, simple geometries such as strips^7,8^, rings^9^, sheets^10^ and meshes^2,9^ are produced. While also balloon-shaped single ventricles have been generated using casting molds^11^ or by seeding cells onto prefabricated, including 3D printed, scaffolds^12-14^, these techniques are ultimately limited in their potential to generate tissues with complex architecture or defined multicellular arrangements.

Recently, first steps have been achieved towards producing anatomically accurate whole-heart or simplified heart models, as well as functional units thereof, utilizing 3D-bioprinting^15-18^. Noor *et al*. employed a patient-specific approach to generate a miniaturized heart comprising chambers and major vessels. Human induced pluripotent stem cell-derived cardiomyocytes (hiPSC-CMs) embedded in an omentum-derived matrix were printed directly. However, no electrical activity or contractility was demonstrated^17^. Lee *et al*. developed and applied freeform reversible embedding of soft hydrogels (FRESH v.2.0) to print a high concentration collagen outer shell into which a fibrin-based gel comprising embryonic stem cell-derived CMs and cardiac fibroblasts was deposited^16^. These fabricated ventricles developed interconnected networks of CMs and showed spontaneous electrical activity and contractions. This “indirect” approach with the necessity of generating a shell structure, however, appears to limit the geometric complexity that can be achieved. Kupfer *et al*. printed simplified heart models using a single bioink containing hiPSCs, which were differentiated into CMs *in situ*^15^. This approach requires a method to locally differentiate hiPSCs in different cell types and raises the risk of teratoma formation.

Given the current limitations of biofabrication, especially in the field of cardiac tissue engineering, we sought to develop a method that allows for direct printing of hiPSC-CMs into functional heart tissue, based on a modified FRESH approach^16,19^. Here, we report a simple and effective strategy to directly 3D-bioprint cardiac tissues ranging from simple rings to centimeter-sized functional models of human cardiac ventricles that can be cultured for at least 100 days and respond to pharmacological stimuli.

## Results

### Simple method to produce a support bath for in-gel printing

Collagen is the most abundant ECM protein in the heart, making it a prime candidate for cardiac tissue engineering. We and others have previously demonstrated the suitability of collagen hydrogels for cardiac tissue engineering^9,20^. While collagen-hydrogels are difficult to print once gelled, pre-gel solutions of collagen can be printed into a support bath (**Fig. 1a**)^16,19^.

**Fig. 1:**
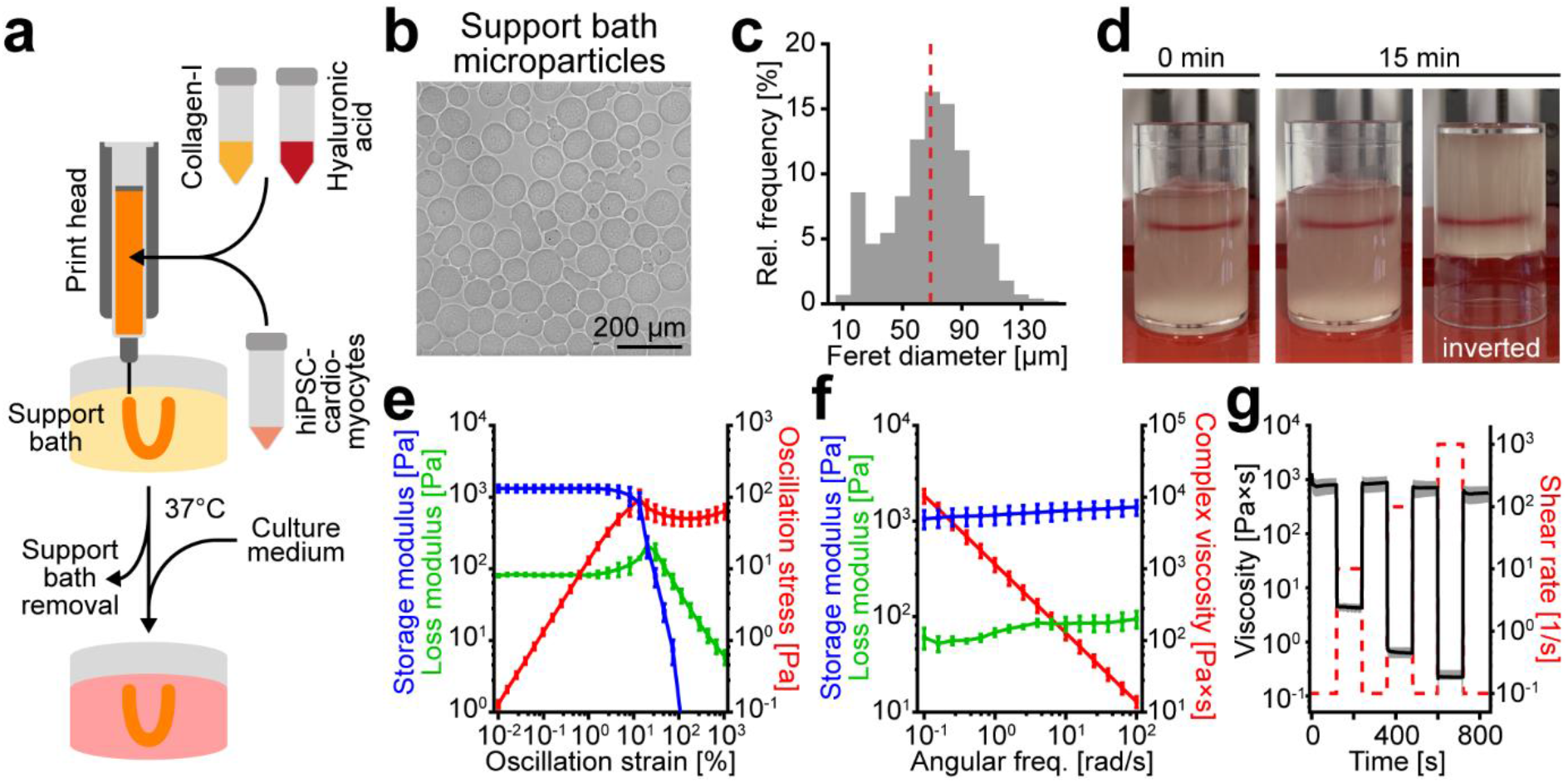
Compacted microparticles display shear thinning and self-healing properties. **a**, Scheme illustrating the in-gel bioprinting approach using collagen-based bioinks and a thermo-responsive microparticle support bath. **b**, Representative phase contrast image of gelatin/gum arabic microparticles produced by complex coacervation. **c**, Size distribution of the microparticles (n = 14 081). Red dotted line: mean Feret diameter. **d**, Images of red food dye printed into compacted microparticles subsequent to printing and 15 min thereafter. **e**, Storage and loss moduli and oscillation stress of the microparticle support bath over an oscillatory amplitude sweep (strain: 0.01-1000%, angular frequency: 10 rad/s; n = 3). **f**, Storage and loss moduli and value of complex viscosity η of the microparticle support bath over an oscillatory frequency sweep (angular freq.: 0.1–100 rad/s, strain: 1%; n = 3). **g**, Viscosity recovery behavior upon alternating shear rates (0.1 s^-1^ *vs*. 10, 100, or 1000 s^-1^) of the microparticle support bath in rotational rheological measurement (n = 3).

This so-called in-gel printing requires a support bath that on one hand provides enough stability to retain the printed liquid ink’s shape and on the other hand allows the printing nozzle to pass through the support bath. In order to print hiPSC-derived cardiomyocytes (hiPSC-CMs) into complex shapes, we developed, based on the work of Lee et al.^16^ (“FRESH v2.0”), a simplified protocol to produce gelatin/gum arabic microparticles by complex coacervation at a scale of two hundred milliliters (**Supplementary Fig. 1a**). Microparticles were predominantly circular in shape, with some exhibiting elongated elliptical shape (diameter: 69 ± 27 μm, data are ± SD, **Fig. 1b,c**). When compacted by centrifugation, microparticles formed a support bath with self-healing properties, permitting a printing nozzle to pass through it freely (**Supplementary Fig. 1b**). Food dye dispensed into the support bath was retained in place, even when the container was inverted (**Fig. 1d**). To corroborate these observations, rheological analyses were performed. The support bath showed rheological solid-like behavior (storage modulus G’ = 1320.7 Pa, loss modulus G’’ = 82.5 Pa) at low strains and started to yield at a critical amplitude of 8.2% strain (Oscillation stress: 71.6 Pa; **Fig. 1e**). A frequency sweep showed solid-like behavior in the entire frequency range tested as well as shear thinning behavior (**Fig. 1f**). Moreover, the support bath exhibited near instantaneous and complete recovery behavior at alternating shear rates (**Fig. 1g**).

Taken together, we have developed a simple method for the production of a support bath with self-healing properties suitable for in-gel printing.

### Collagen/hyaluronic acid ink enables accurate and reproducible printing of complex structures

3D-bioprinting promises to generate tissue constructs reproducibly and accurately, based on a given 3D model. To assess the printability of collagen in the above described support bath, acellular ring-shaped constructs were printed with collagen-inks at 1.0, 2.0, or 3.0 mg/ml. Notably, collagen is commonly dissolved in acid for long term storage and thus has to be neutralized before incorporating cells. As neutralization initiates gel formation, which happens faster at higher temperatures, the collagen-ink was prepared on ice and the printhead was pre-cooled (10-15°C). To determine the uniformity of printed rings and thus reproducibility of the printing procedure, overlap-maps were generated from images of multiple printed rings (**Fig. 2a**). With increasing collagen content, overlap-maps showed more uniform printed rings with smoother borders. To increase the reproducibility, hyaluronic acid (HA, 1.5 and 3.0 mg/ml), a glycosaminoglycan found in the ECM, which has previously been described to improve the printability of bioinks^21^, was added to the collagen. HA especially improved the uniformity of rings printed at 2.0 and 3.0 mg/ml collagen. These findings were further corroborated by radial intensity profiles (**Fig. 2b**).

**Fig. 2:**
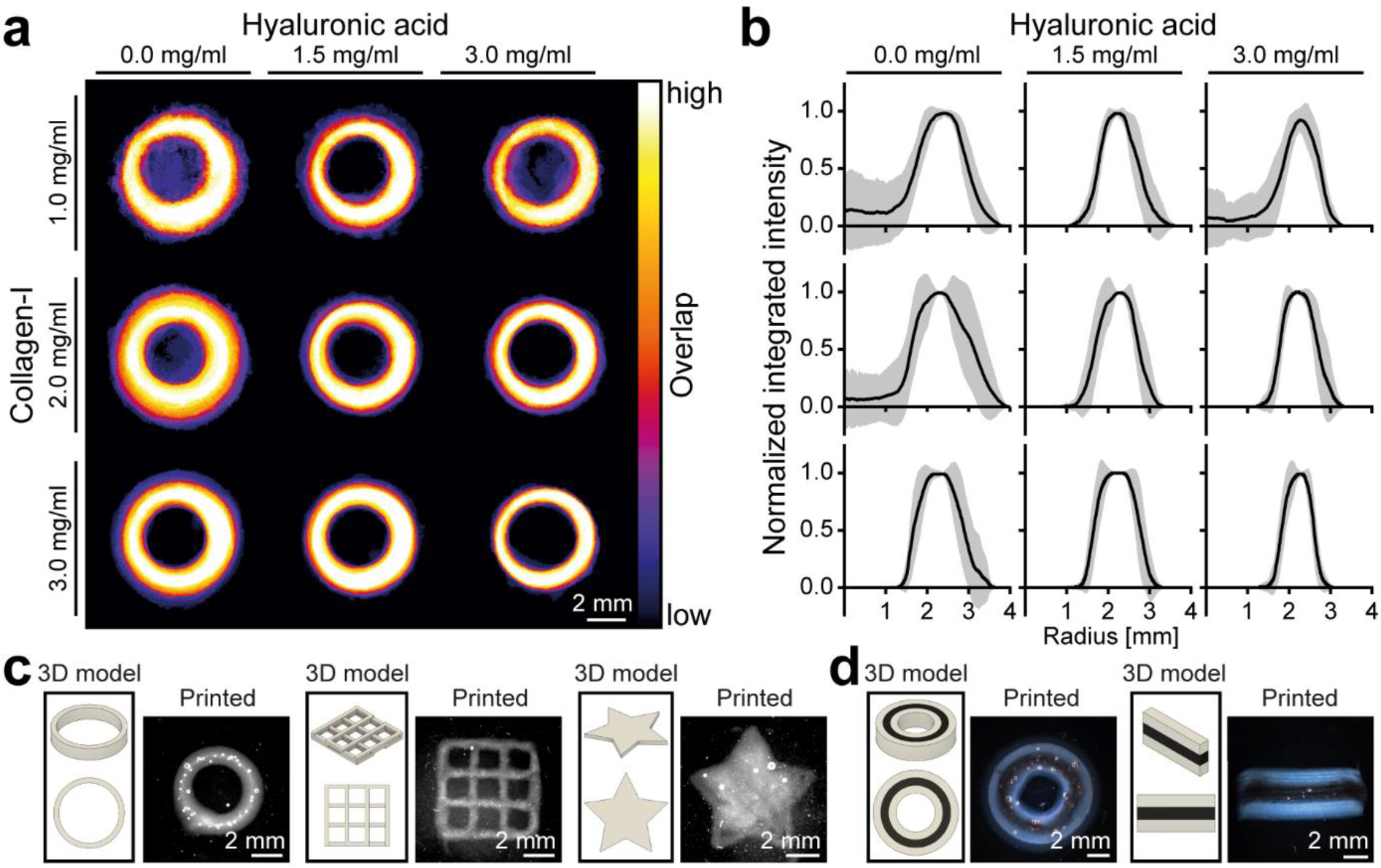
Collagen/hyaluronic acid ink enables printing of stable structures. **a**, Overlap-maps of darkfield images of rings printed with inks comprising 1.0, 2.0, or, 3.0 mg/ml collagen-I and optionally 1.5 or 3.0 mg/ml hyaluronic acid (n = 14 to 23). **b**, Radial intensity profiles (data are mean ± SD) of rings printed in (**a**) (n = 14 to 23). **c**, 3D models of ring-, grid- and star-shaped geometries and representative darkfield images of resultant constructs printed using an ink comprising 3.0 mg/ml collagen-I and 3.0 mg/ml hyaluronic acid (*CollHA3-3*). **d**, 3D models of multi-material geometries (concentric rings, stacked rectangles) and representative darkfield images of resultant constructs printed using *CollHA3-3* with and without a contrast agent.

With increasing collagen concentrations and the addition of HA, intensity profiles showed less variance and became more well-defined. The bioink comprising 3.0 mg/ml collagen-I and 3.0 mg/ml HA (*CollHA3-3*) was selected for further studies, as it offered the highest reproducibility and best control over extrusion speed in our technical setup, as indicated by the thinner wall thickness of those rings.

To demonstrate accuracy of the printing process, different geometries (ring-, grid-, and star-shaped) were printed. The printed constructs showed well-defined sharp edges and were found to accurately reproduce the underlying 3D models (**Fig. 2c**). Printing *CollHA3-3* with and without a contrast agent alternatingly (concentric rings or stacked rectangles) resulted in continuous and stable constructs, demonstrating that the two bioinks had fused both within a layer and across layers (**Fig. 2d**).

Collectively, these findings show that neutralized collagen-based inks can be printed accurately to form stable multi-layered constructs and thus has the potential to enable printing of complex tissues.

### Reproducible and accurate 3D-bioprinting of ring-shaped functional cardiac tissues

To generate a hierarchical structured tissue or organ, it is important to have control over cell distribution. Therefore, cells should be incorporated into the ink itself before printing. To assess whether the addition of cells influences the ink’s printability, we incorporated 25 Mio/ml hiPSC-CMs into the *CollHA3-3* ink to print ring-shaped tissues (specified height: 1 mm (five-layered), specified diameter: 5 mm). The resultant rings were stable and showed good uniformity, as illustrated by an overlap-map (**Fig. 3a**) and further supported by a radial intensity profile (**Supplementary Fig. 2a**), indicating that the addition of cells did not have a detrimental effect on printability.

**Fig. 3:**
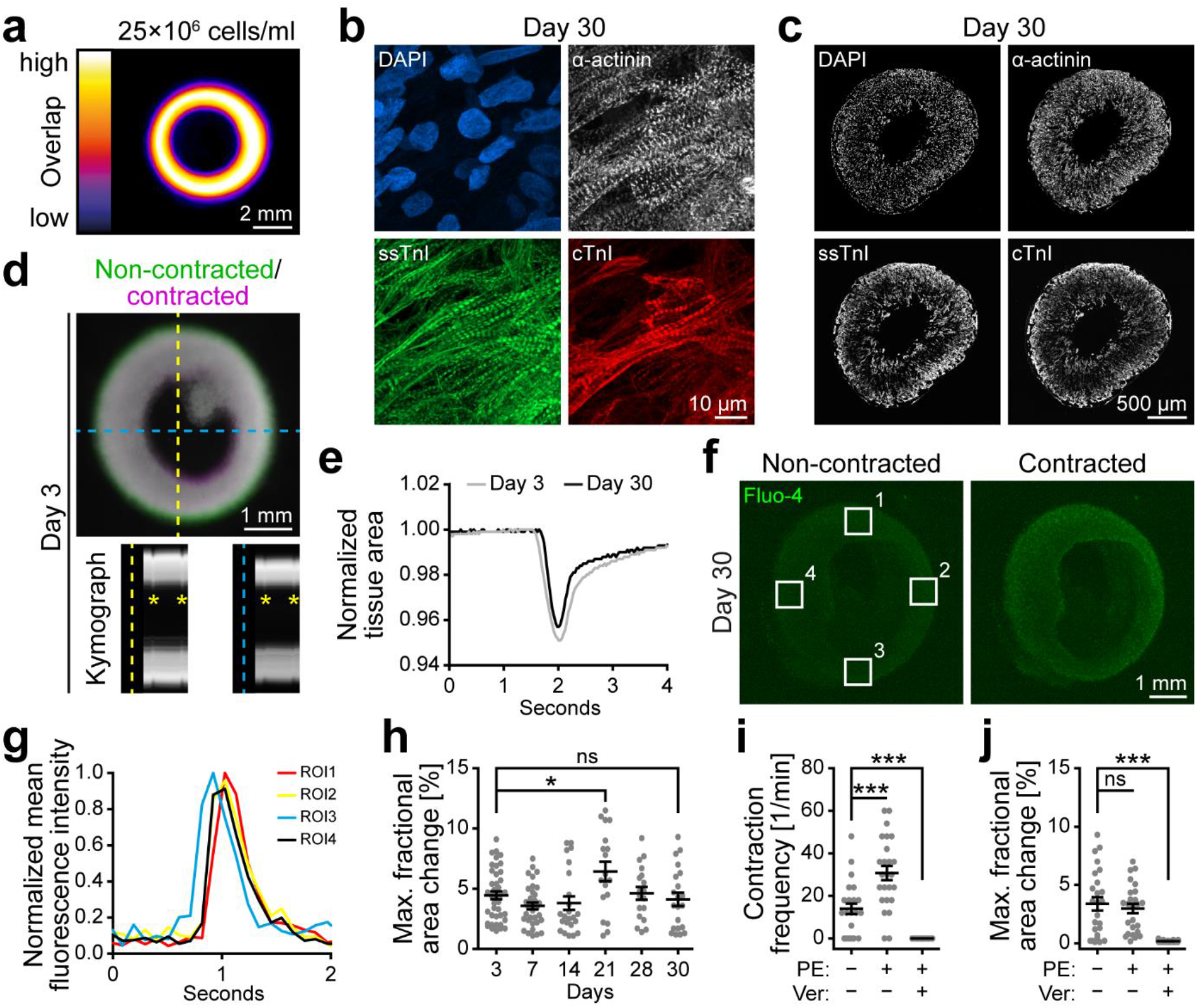
Printed hiPSC-derived cardiomyocytes form functional cardiac tissues. **a**, Overlap-map of rings printed with *CollHA3-3* containing 25 Mio/ml hiPSC-CMs (n = 119). **b,c**, Representative maximum intensity projections of confocal z-stacks of wholemount-stained (**b**) and images of stained cryosections (**c**) of printed tissues after 30 days of culture. Sarcomeric proteins: α-actinin, slow skeletal troponin I (ssTnI), cardiac troponin I (cTnI). DNA: DAPI. **d**, Visualization of contraction. Top: merged false-colored darkfield images of a non-contracted (green) and contracted (magenta) printed ring on day 3. Bottom: Kymograph analyses along vertical (yellow) and horizontal (blue) axis. **e**, Representative plots of normalized projected tissue area over the course of a single contraction at days 3 and 30. **f,g**, Representative images of calcium flux in non-contracted and contracted printed tissue (day 30) visualized with Fluo-4 (**f**) and its quantification (**g**) based on normalized mean fluorescence intensity over the course of contraction in: regions of interest (ROI1–4, white boxes) in (**f**). **h**, Maximum fractional change in projected tissue area during contraction of printed rings over the course of cultivation from day 3 to 30 (n = 18 to 42). **i**, Contraction frequency of printed rings (day 30) at baseline and after treatment with phenylephrine (PE, 50 μM) and verapamil (Ver, 1 μM; n = 25). **j**, Maximum fractional change in projected tissue area during contraction of printed rings (day 30) at baseline and after treatment with PE (50 μM) and Ver (1 μM; n = 25). Statistics: ANOVA followed by post analysis according to Dunnett. Data are mean ± SEM. *: p < 0.05. ***: p < 0.001. ns: not significant.

To determine if 3D printing affects cell survival, printed tissues and such produced by drop-casting, were stained using calcein-AM/ethidium homodimer-1 (EthD-1) immediately after tissue construction (day 0) and 7 days post fabrication (day 7) (**Supplementary Fig. 2b**). In printed and cast tissues, hiPSC-CMs were predominantly viable (calcein-positive) at day 0 and day 7 and more elongated and spread out at day 7 compared to day 0 (**Supplementary Fig. 2b**). No significant difference in viability was found between cast and printed tissues at either timepoint (**Supplementary Fig. 2c**).

In addition to viability, we investigated the morphology and distribution of hiPSC-CMs by immunofluorescence staining of sections as well as whole tissues at day 30. Printed hiPSC-CMs showed typical striated patterns of sarcomeric α-actinin as well as slow skeletal troponin-I (ssTnI) and cardiac troponin-I (cTnI) (**Fig. 3b**). While α-actinin and ssTnI were expressed at similar levels by most cells, cTnI, a marker of more mature hiPSC-CMs, was more varied across neighboring cells. Notably, hiPSC-CMs were present throughout the printed tissues as evidenced by nuclear staining and had formed interconnected networks, as demonstrated by evenly distributed expression of α-actinin (**Fig. 3c**). In contrast, expression levels of ssTnI increased towards the edges of the tissues. This differential expression pattern was even more pronounced for cTnI. These findings show that printed hiPSC-CMs established organized sarcomeres and that the printed tissues comprise interconnected hiPSC-CMs of different states of maturation.

Previous studies printing hiPSC-CMs in a direct manner have not shown contractility of the resultant tissues^17,18^. Video microscopy of 57 samples revealed that 45 of our printed tissues exhibit spontaneous, concentric contractions already at day 3 (**Fig. 3d** and **Supplementary Video S1**) and only one of the 57 rings did not initiate contraction during the observation time. Contractions were associated with a transient decrease in projected tissue area (**Fig. 3e**). Moreover, printed tissues displayed a synchronized influx and efflux of calcium over the course of a contraction cycle as evident by increasing and decreasing Fluo-4 fluorescence intensity (**Fig. 3f** and **Supplementary Video S2**). Measuring the fluorescence intensity at different regions allowed the identification of a pacing center (ROI3, **Fig. 3g**) and showed high synchronicity over time (**Supplementary Fig. 2d**). Distant regions reached peak fluorescence intensity with a delay of ∼100 ms (1 frame) compared to the pacing center (**Fig. 3g**). These observations are further evidence for the formation of interconnected networks of hiPSC-CMs within the printed tissues. While spontaneous contraction frequencies did not significantly change between day 3 and day 30, we observed a transient reduction in frequency between day 7 and day 21 (**Supplementary Fig. 2e**). As printed tissues were cultured free floating, accurate force measurements could not be performed. Instead, the change in projected tissue area during a contraction cycle was used as a surrogate marker for contraction amplitude (**Fig. 3d,h**). Notably, maximum fractional area change was similar from day 3 to day 30.

To be suitable as model systems or transplants, engineered cardiac tissues must show physiological responses to pharmacological stimulation. Printed tissues were thus treated first with adrenergic agonist phenylephrine (PE), followed by verapamil (Ver), an inhibitor of L-type calcium channels (**Supplementary Fig. 2f**). PE-stimulation resulted in a doubling of contraction frequency (30.7 ± 3.3 min^-1^ *vs*. 13.9 ± 2.4 min^-1^, data are mean ± SEM), but no significant change in the maximum fractional area change (**Fig. 3i,j** and **Supplementary Video S3**). Subsequent addition of verapamil stopped the contractions (**Fig. 3i,j** and **Supplementary Video S3**).

Taken together, we show that the procedure and bioink developed here enable printing of hiPSC-CMs into functional cardiac tissue that is responsive to pharmacological stimulation.

### Printed cardiac tissues exert concentric forces against passive resistance

To determine whether the printed rings can exert concentric forces similar to the heart, we designed and fabricated an array of six flexible pillars (**Fig. 4a**). Printed rings were transferred onto the pillar array on day 1 and cultured until day 30. During this time, they compacted around the pillars (**Fig. 4b**). Notably, printed rings were able to synchronously deflect all 6 pillars during a contraction cycle, illustrating their ability to exert a force against resistance (**Supplementary Video S4**).

**Fig. 4:**
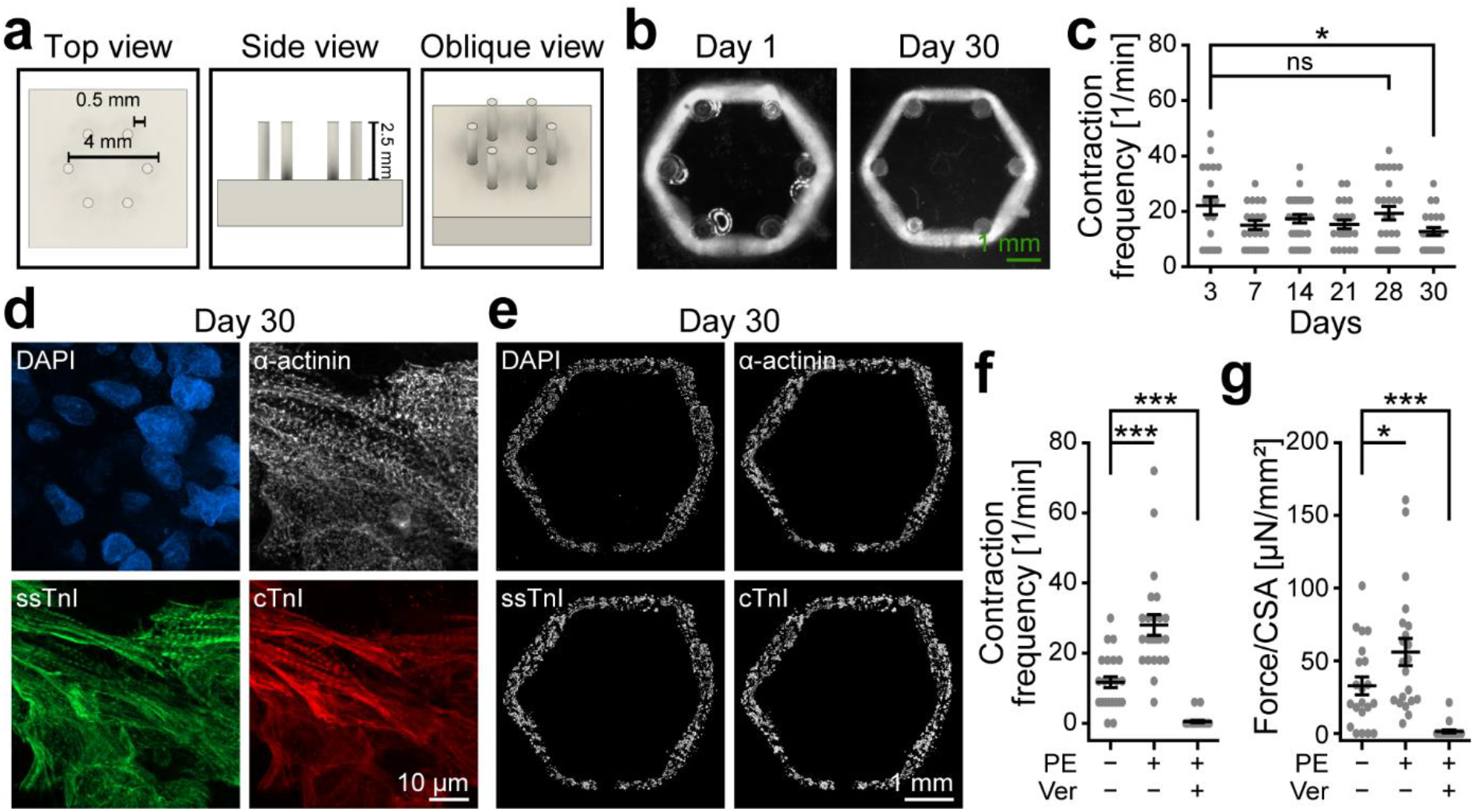
Printed cardiac tissues exert concentric forces against passive resistance. **a**, 3D model of six pillar array. **b**, Representative darkfield images of printed rings cultured on flexible pillar arrays on day 1 and 30. **c**, Contraction frequency of printed rings cultured on flexible pillar array over the course of cultivation from day 3 to 30 (n = 19–29; P < 0.05 vs. day 3 by ANOVA followed by post analysis according to Dunnett). **d,e**, Representative maximum intensity projections of confocal z-stacks of wholemount-stained printed tissues (**d**) and stained cryosections of printed tissues (**e**) cultured on flexible pillar array (day 30). Sarcomeric proteins: α-actinin, slow skeletal troponin I (ssTnI), cardiac troponin I (cTnI). DNA: DAPI. **f**, Contraction frequency of printed rings (day 30) at baseline and after treatment with phenylephrine (PE, 50 μM) and verapamil (Ver, 1 μM; n = 24. **g**, Force exerted by printed rings cultured on flexible pillar array (day 30) normalized to tissue cross sectional area (CSA) at baseline and after treatment with phenylephrine (PE, 50 μM) and verapamil (Ver, 1 μM; n = 21). Statistics: ANOVA followed by post analysis according to Dunnett. Data are mean ± SEM. *: p < 0.05. ***: p < 0.001. ns: not significant.

Like their free-floating counterparts, the rings showed spontaneous contractions as early as day 3 (day 3: 19 of 38, day 7: 29 of 38). Only one of the 38 printed rings did not initiate contraction until day 30 but initiated contractions upon PE-stimulation. Visualizing calcium fluxes using Fluo-4 indicated highly synchronized tissues (**Supplementary Fig. 3a,b** and **Supplementary Video S5**). Only in one of the five analyzed rings small clusters of cells were observed that showed additional “out of rhythm” beats (**Supplementary Fig. 3a,b** and **Supplementary Video S6**). Contraction frequency remained stable at around 20 beats per minute up to day 28 and was slightly reduced at day 30 (**Fig. 4c**). Immunofluorescence staining of whole tissues (**Fig. 4d**) or sections (**Fig. 4e**) for sarcomeric markers showed that hiPSC-CMs exhibited typical striated patterns of α-actinin, ssTnI, and cTnI at day 30 (**Fig. 4d**). Similar to free-floating rings, α-actinin and ssTnI were expressed at similar levels by most hiPSC-CMs, while cTnI was more varied across neighboring cells. However, in contrast to free-floating rings, expression of ssTnI and cTnI was found to be evenly distributed throughout the tissue in printed rings working against passive resistance (**Fig. 4e**).

To determine if printed rings react to pharmacological stimulation also under passive mechanical stimulation, PE- and Ver-treatment was performed at day 30 (**Supplementary Video S7)**. PE-stimulation significantly increased contraction frequency in rings under passive mechanical stimulation, similar to free-floating rings (**Fig. 4f**). Forces exerted by the printed tissues were calculated based on pillar deflection (**Supplementary Fig. 3c,d**) and showed an increase upon PE-stimulation (**Fig. 4g**). Ver-treatment resulted in significantly reduced contraction frequency and force generation, whereby 22 out of 24 analyzed rings stopped beating (**Fig. 4f,g**).

Taken together, we introduce a new six-pillar system to culture ring-shaped tissues and show that printed cardiac tissues containing hiPSC-CMs can exert concentric forces against passive resistance.

### Collagen/hyaluronic acid ink enables 3D-bioprinting of a functional model of a human heart ventricle

The goal of 3D-bioprinting is to enable the fabrication of complex hierarchical structures. To test whether our approach enables 3D-bioprinting of a functional model of a human heart ventricle, a model 14 mm in height and 8 mm in diameter at its widest part was designed and printed (**Fig. 5a**).

**Fig. 5:**
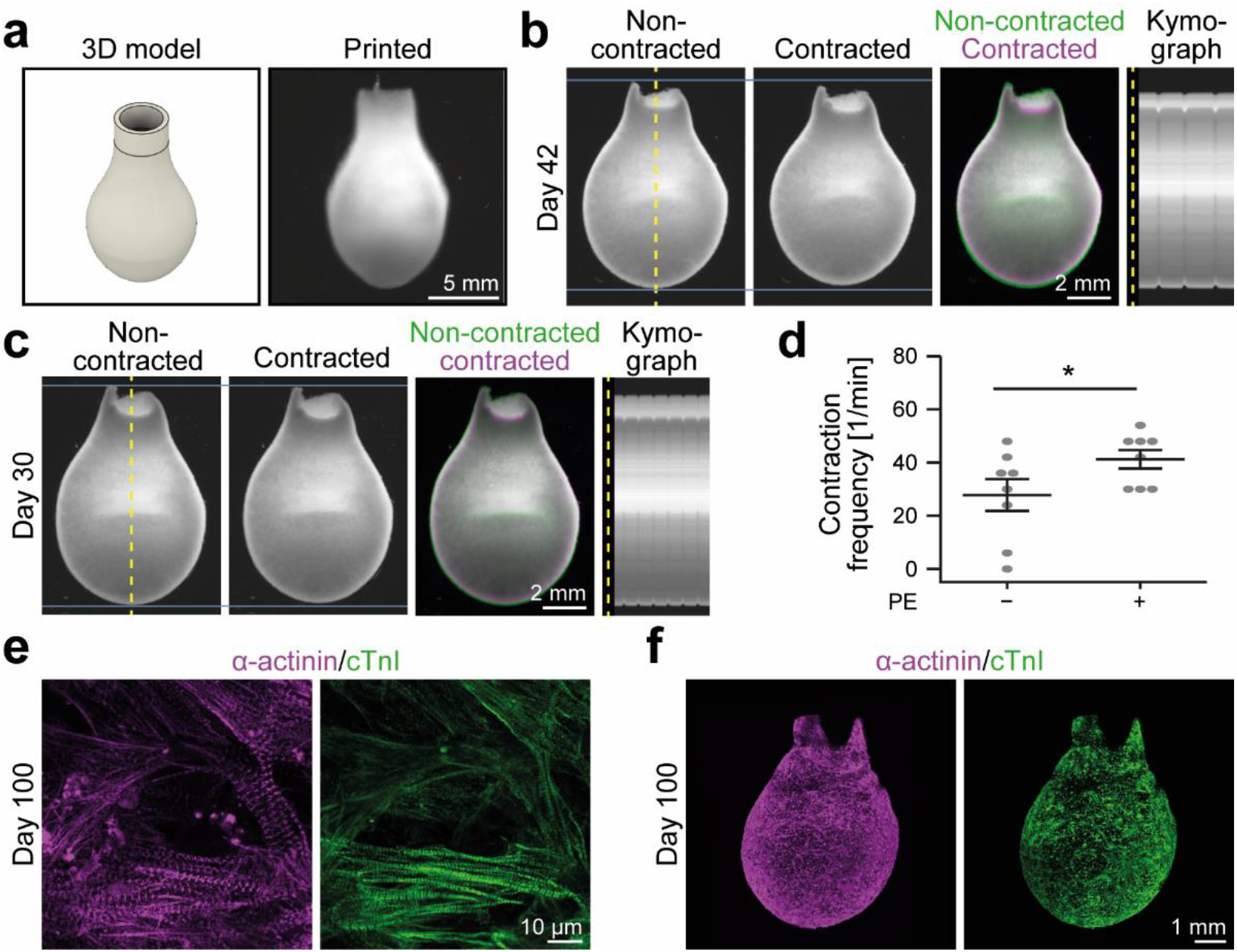
In gel approach enables 3D-bioprinting of functional ventricle models. **a**, 3D model of a ventricle and representative darkfield image of a printed ventricle. **b, c**, Visualization of contraction of two printed ventricles at day 42 (**b**) and day 30 (**c**). Left: representative individual darkfield images of a non-contracted and contracted printed ventricle. Middle: merged false-colored images (non-contracted: green; contracted: magenta). Right: Kymograph analyses along vertical axis (yellow dotted lines). **d**, Contraction frequency of printed ventricles at day 30 before and after treatment with phenylephrine (PE, 50 μM; n = 8; *: p < 0.05, paired t-test). **e,f**, Detail view (confocal, **e**) and overview maximum intensity projections of z-stacks of whole printed ventricles at day 100. For overview images acquired in LSFM (**f**), samples were optically cleared. Sarcomeric proteins: α-actinin and cardiac troponin I (cTnI).

These ventricles were stable and developed spontaneous synchronous contractions as early as 7 days after fabrication, which persisted over the course of cultivation for up to 100 days (**Fig. 5b,c** and **Supplementary Video S8,9**). Moreover, PE-stimulation induced an increase in contraction frequency of printed ventricle models cultured for 30 days from 27.8 ± 6.0 to 41.3 ± 3.5 beats per minute (p < 0.05, **Fig. 5d** and **Supplementary Video S10**). Finally, analysis of stained ventricles revealed that hiPSC-CMs exhibited typical striated patterns of α-actinin and cTnI at day 100 (**Fig. 5e**). In optically cleared samples imaged by light-sheet fluorescence microscopy (LSFM), hiPSC-CMs were found to be distributed over the whole printed tissue, with both α-actinin and cTnI showing a similar expression pattern (**Fig. 5f**).

Taken together, we demonstrate direct 3D-bioprinting of centimeter-sized functional models of human cardiac ventricles that can be cultured for at least 100 days.

## Discussion

Our findings demonstrate that it is possible to directly print hiPSC-CMs into functional cardiac tissues. Several lines of evidence support this conclusion. hiPSC-CMs embedded in a collagen/hyaluronic acid ink could be accurately and reproducibly 3D-bioprinted into ring- and ventricle-shaped cardiac tissues, in which hiPSC-CMs formed interconnected networks. Concurrently, printed tissues exhibited spontaneous and synchronous contractions that persisted for several months. These contractions could be modulated in frequency by pharmacological stimulation. Furthermore, printed tissues were able to contract against passive resistance, which was associated with compaction and maturation.

Similar to previously described biofabricated heart ventricles^11-13,15^, the forces generated by the here printed constructs are far below forces generated by cast tissues or the adult human heart^7-9,22^. Thus, to generate cardiac ventricles with physiologically relevant pump function, it is important to apply approaches utilized in cast tissues to enhance in the future contractility, such as incorporation of other cell types (e.g. fibroblasts^9^), electroconductive materials^20,23,24^, and pro-proliferative factors^25^, as well as electrical and mechanical stimulation^7^ or metabolic maturation^26,27^. In addition, new protocols are needed to enhance the maturation of hiPSC-CMs.

While there are still many challenges to overcome, the here developed method opens up new possibilities for generating complex functional heart tissues. The possibility to directly print hiPSC-CMs together with printing other cell types such as fibroblasts and vascular cells, will allow printing healthy, as well as diseased heart models, containing, besides myocardium, for example major vessels, the fibrous skeleton of the heart, valves, and/or scarred areas. Furthermore, the ability to precisely place different cardiomyocyte subtypes (i.e. atrial, ventricular or pace-maker cardiomyocytes) plus the addition of electroconductive materials to the bioink may further allow in the future for guiding action potential propagation throughout the tissue to more accurately replicate cardiac electrophysiology.

Taken together, our method provides a step towards the fabrication of advanced drug screening models, tissue grafts, and in the future maybe even an engineered heart for organ transplantation.

## Methods

### Support bath preparation

Equal volumes of aqueous solutions of 4% (w/v) gelatin type A (Sigma Aldrich) and 4% (w/v) gum arabic (Carl Roth, solution neutralized to pH 7.0) were combined at 50-55°C under constant stirring (600 rpm; **Supplementary Fig. 1a**). To induce microparticle formation, the pH was lowered to 3.8-4.0 using hydrochloric acid (HCl). While maintaining stirring (600 rpm), microparticles were rapidly cooled by adding an equal volume of ice cold (<4°C) ultrapure water to the microparticle preparation and placing ice around the beaker until a temperature of ∼15°C was reached. The microparticles were then left to settle at 4°C, decanted, washed with ultrapure water at room temperature and collected in centrifugation tubes. Subsequent steps were performed at room temperature. Microparticles were washed six times with sterile DPBS. During washing steps, microparticles were compacted by centrifugation for 3 min at 300 x g, rid of supernatant, and subsequently resuspended in fresh DPBS. After the final washing step, compacted microparticles were centrifuged again (3 min, 300 x g) and finally transferred to appropriate vessels using a 1000 μl micropipette. For printing cell-laden bioinks, the final two washing steps were performed with RPMI 1640 (Thermo Fisher Scientific or Sigma-Aldrich). Microparticle Feret diameter was measured in phase-contrast images using a custom designed FIJI^28^ Macro.

### Rheology

Rheological measurements were performed using an AR-G2 rheometer (TA Instruments) equipped with a cone-plate geometry (diameter: 40 mm, gap: 55 μm). Support bath (0.3 ml) was transferred to the baseplate of the rheometer, followed by lowering the top plate to 65 μm. Excess material was removed before lowering the rheometer head to the measurement gap to ensure a suitably filled gap. Measurements of microparticle support bath were performed at 21°C, controlled by the attached Peltier-element. Frequency sweeps were performed at a set 1% strain, amplitude sweeps at set 10 rad/s angular frequency.

### Stem cell culture, cardiac differentiation, and cardiomyocyte expansion

hiPSCs were cultured in StemMACS iPS brew XF (Miltenyi Biotec) on Matrigel- (Corning) or Geltrex- (Thermo Fisher Scientific) coated culture ware. Medium changes were performed daily and hiPSCs were passaged every 2-4 days. For cardiac differentiation, hiPSCs grown to 80-90% confluency were induced to differentiate by changing the medium to differentiation medium, comprised of RPMI 1640 supplemented with 2% B-27 Minus Insulin (both Thermo Fisher Scientific) and 100 μM ascorbic acid. For the first 24 h, medium was supplemented with 8-10 μM CHIR99021 (Hycultec), followed by 48 h without additional growth factors and 48 h of treatment with IWR1-endo (Selleckchem) ^29^. Differentiated hiPSC-CMs were maintained in RPMI 1640 supplemented with 2% B-27 (Thermo Fisher Scientific) from day 7 on, then purified by metabolic selection in RPMI 1640 without glucose (Thermo Fisher Scientific), supplemented with 5 mM sodium-DL-lactate and 100 μM ascorbic acid for 5 days. Subsequently, purified hiPSC-CM were dissociated, replated at low density (25.000-30.000 cells/cm^2^) and expanded using 3 μM CHIR99021 for 5-8 days as previously described by Buikema and colleagues^30^.

### Bioink preparation and bioprinting

All steps were performed at room temperature if not stated otherwise. HA (1.5-1.8 MDa, Sigma-Aldrich) was dissolved in Iscove’s Modified Eagle’s Medium (IMDM, Thermo Fisher Scientific) to produce a 2% (w/v) solution. HA was combined on ice with culture medium and an appropriate amount of neutralization solution (1/9 of collagen-I solution; Advanced Biomatrix), before acid-solubilized rat tail collagen-I (3.9-4.1 mg/ml; Advanced Biomatrix) was added. For bioinks, cells were centrifuged (300 x g, 3 min), rid of supernatant and resuspended in the ink by careful pipetting. The bioink was then transferred to a 1 ml syringe. A custom designed cooling sheath was placed around the syringe and the assembly was mounted into the printhead of a commercial bioprinter (Allevi 2), modified to accommodate 1 ml syringes. Pre-cooling of the print head and cooling sheath allowed for temperatures within the print head to be maintained at 13-19°C during printing. Bioinks comprised of 3 mg/ml collagen and 3 mg/ml HA were printed at 4.0-5.5 psi (28-38 kPa) using 23 Ga stainless steel nozzles. Printed constructs were incubated at 37°C for 45 min, after which the support bath was removed by washing with sterile DPBS.

### Cardiac tissue cultivation

Printed and DPBS-washed constructs were cultured in medium comprised of Iscove’s Modified Eagle’s Medium (IMDM), supplemented with 4% B-27 Minus Insulin, 1 mM non-essential amino acids (NEAA, all Thermo Fisher Scientific), 0.1 μg/ml IGF-1, 10 ng/ml FGF-2, 5 ng/ml VEGF-165 (all PeproTech), and 100 U/ml Penicillin/Streptomycin (Thermo Fisher Scientific) (based on Tiburcy *et al*.)^9^. For the first 24 h after printing, culture medium was supplemented with 10 μM ROCK-inhibitor (Y-27632, Selleckchem). Printed rings were either cultured free-floating or transferred to an array of flexible pillars one day after printing. Culture medium was exchanged every 2 to 3 days.

### Analysis of printing reproducibility

Darkfield images of printed constructs were acquired on a stereoscope (ZEISS Axiozoom). To assess printing reproducibility, a custom designed semi-automatic FIJI Macro was developed. An intensity threshold was applied to images to produce binary masks. The objects within the masks corresponding to the printed rings were isolated and aligned using their inner and outer outlines as reference. A sum intensity projection of all masks was generated and pixel intensities were color coded using a “Fire” lookup table (FIJI). For quantification, radial intensity profiles of individual masks were generated using the “Radial Profile Extended” Plugin for FIJI and averaged.

### Analysis of cell viability

To assess cell viability, printed constructs were incubated with 2 μM EthD-1, 1 μM calcein-AM and 5 μg/ml Hoechst 33342 in DPBS at 37°C for 30 min. Image stacks of 100 μm thickness were acquired on a LSM900 confocal microscope (Zeiss), equipped with a Plan-Apochromat 20x/0.8 M27 objective (Zeiss). To identify the proportion of viable cells, a manually set intensity threshold was applied to maximum intensity projections of image stacks. Subsequently, using ImageJ’s ‘Analyze Particles’ tool, calcein- and EthD-1-positive cells were identified. Percentage viability was calculated from the number of calcein-positive cells divided by the total number of cells (calcein-positive cells + EthD-1-positive cells).

### Cryoembedding and sectioning

Printed ring- or ventricle-shaped tissues were fixed with 4% formalin at room temperature for 30 min or 2 h, respectively. Subsequently, samples were washed with PBS and incubated in 15% sucrose and then 30% sucrose (w/v, in PBS), both until samples sank to the bottom of the reaction tube. Samples were embedded in Tissue-Tek O.C.T. Compound (Sakura Finetek) and frozen in vapor phase liquid nitrogen. Sections of 7 μM thickness were obtained using a cryostat (LEICA), mounted onto glass slides, and stored at -80°C.

### Immunofluorescence staining

Cryosections were washed twice with PBS, permeabilized with 0.5% Triton-X100/PBS for 10 min, and incubated in blocking buffer (0.2% Tween-20, 5% BSA in PBS) for 30 min. Sections were incubated with primary antibodies in blocking buffer overnight at 4°C using the following antibodies (all Abcam): anti-sarcomeric-alpha-actinin (ab9465; 1:500), anti-cTnI (ab56357; 1:500), and anti-ssTnI (ab203515; 1:250).

Sections were then washed three times with 0.1% NP-40/PBS and incubated with secondary antibodies in blocking buffer for 2 h at room temperature. The following secondary antibodies were used (all Thermo Fisher): anti-mouse Alexa Fluor 647 (1:500), anti-goat Alexa Fluor 594 (1:500), and anti-rabbit Alexa Fluor 488 (1:500). Next, sections were washed twice with 0.1% NP-40/PBS, counterstained with DAPI (0.5 μg/ml in 0.1% NP-40/PBS for 15 min and washed with PBS. For whole mount staining of printed tissues, samples were permeabilized in 0.5% Triton-X100 for 30 min, blocked for 1 h, incubated in primary antibody solution overnight for ring-shaped or 48 h ventricle-shaped tissues, washed three times with 0.5% NP-40/PBS and incubated in secondary antibody solution for 2 h at room temperature for ring-shaped or overnight at 4°C for ventricle-shaped tissues. Printed rings were counterstained with DAPI (0.5 μg/ml in 0.1% NP-40/PBS for 30 min. Cover glasses were mounted onto tissue samples using Fluoromount G. 3D-printed plastic spacers (1 mm thickness) were used to preserve the 3D structure of whole-mount tissues.

### Confocal and Light-Sheet Fluorescence Imaging (LSFM)

Images of cryosections and ring-shaped constructs were acquired using an LSM900 confocal microscope (Zeiss), equipped with Plan-Apochromat 20x/0.8 M27, W Plan-Apochromat 40x/1.0 DIC M27 and Plan-Apochromat 63x/1.4 Oil DIC M27 objectives (all Zeiss). For imaging whole printed ventricles, optical clearing was performed using a modified ECi protocol.^31^ For LSFM; stained samples were dehydrated in ascending alcohol series (30%, 50%, 70%, 100%, 100%) for 8–12 h at 4°C. For optical clearing, ventricles were incubated in ECi (catalog no. 112372; Sigma-Aldrich, Darmstadt, Germany) for 12 h, and stored in freshly prepared ECi solution at room temperature for another 12 h before imaging in a custom-built light sheet fluorescence microscope. Major parts of the LSFM setup have been described before^32^. Here, laser lines of 488, 561, and 640 nm (OBIS; Coherent Corp., Santa Clara, CA, USA) were used for fluorescence excitation via a customized fiber-coupled laser combiner and a telescope (BE03MA, Thorlabs, Newton, NJ, USA) for beam diameter adjustment. For image processing, the image background from camera noise was subtracted from all tiles and channels. Next, multicolor LSFM image tiles of ventricles were stitched by FIJI^28^ using the BigStitcher^33^. Fused images were exported as 16 bit integer tif-files while preserving the original data resolution (voxel size 2.6×2.6×5.0 μm). Finally, the data was converted into the Imaris file format (Imaris 10.0.0, Bitplane AG, Zurich, Switzerland) for visualization.

### Analysis of contractility

Darkfield videos of printed tissues were recorded using a stereoscope (ZEISS Axiozoom) at 33 frames per second (fps). Contraction frequencies were extrapolated from 10 s videos. To assess fractional area changes, an intensity threshold was applied to videos and the area of the tissue was recorded over time. Area measurements were normalized to each tissue’s maximum area over the course of contractions. For analysis of drug response, subsequent to baseline recording, phenylephrine (PE, 50 μM) was added to the culture medium and videos were recorded after incubation for 10 min at 37°C. Subsequently, verapamil (Ver, 1 μM) was added to the culture medium and the tissues incubated again for 10 min at 37°C before recording videos.

### Analysis of calcium flux

To analyze calcium flux in printed tissues, Fluo-4 reagent solution (prepared according to manufacturer’s instructions; Fluo-4 Direct Calcium-Assay-Kit, Thermo Fisher Scientific) was added to culture medium (50% of culture medium volume) and tissues were incubated at 37°C for 30 min. Subsequently, medium was replaced with fresh culture medium supplemented with PE (50 μM). Tissues were incubated for 10 min at 37°C and fluorescence videos were recorded using a stereoscope (ZEISS Axiozoom) at a framerate of 10 fps.

### Force measurements in printed rings

Flexible pillar arrays were generated using a custom designed mold machined from polyoxymethylene (Supplemental Figure 4A). Sylgard 184 silicone elastomer (DOW chemicals) was prepared at a ratio of 14:1, injected into the mold, degassed and cured at 65°C overnight. Contractile forces *F* were calculated as previously described^34^, according to Hooke’s law:

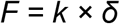

with pillar deflection *δ* and spring constant *k* determined at the height of the tissue above the pillar base (*h*_*tissue*_). For improved spatial resolution, pillar deflection was measured at the pillar top (*δ*_*top*_) with images acquired using a stereoscope (ZEISS Axiozoom). The height of the tissue above the pillar base *h*_*tissue*_ was determined, by focusing the stereoscope on the tissue as well as on the pillar base. Using the Euler-Bernoulli beam theory, the deflection at the tissue height (*δ*_*tissue*_) was calculated as

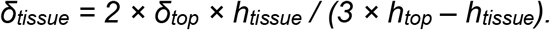

The spring constant *k* was derived from the force-bending relationship at different heights of the pillars as previously described^34^.

## Acknowledgements

We would like to thank the Core Unit Fluorescence Imaging at the Rudolf Virchow Center of the JMU, especially Katherina Hemmen, for technical support with image visualization. This research was funded by the Deutsche Forschungsgemeinschaft [DFG, German Research Foundation, grant numbers: INST 410/91-1 FUGG to F. B. E. and Projektnummer 326998133 – TRR 225 (subprojects C01 to F.B.E.; Z02 to K. G. H. and J. S., A07 to D. W. S., and A01/C02 to B. F.), the Manfred Roth Stiftung (to F. B. E.), and the Research Foundation Medicine at the University Clinic Erlangen, Germany (to F. B. E.).

## Author contributions

Conceptualization: T. U. E. and F. B. E.; Methodology: D. K. and B. F.; K.G. H. and J. S. (tissue clearing and LSFM image processing); Formal analysis: T. U. E., D. K., and B. F.; Investigation: T. U. E., A. A., K. A. M., D. K., S. S., and J. S.; Writing - Original Draft: T. U. E. and F. B. E.; Writing - Review & Editing: all authors; Visualization: T. U. E. and F. B. E.; Supervision: K. G. H., D. W. S., B. F., and F. B. E.; Project administration: F. B. E.; Funding acquisition: K. G. H., D. W. S., B. F., and F. B. E.; all authors read the manuscript and agree with its contents.

