## Supplementary Figure S1 to S3 for "Direct 3D-bioprinting of hiPSC-derived cardiomyocytes to generate functional cardiac tissues"

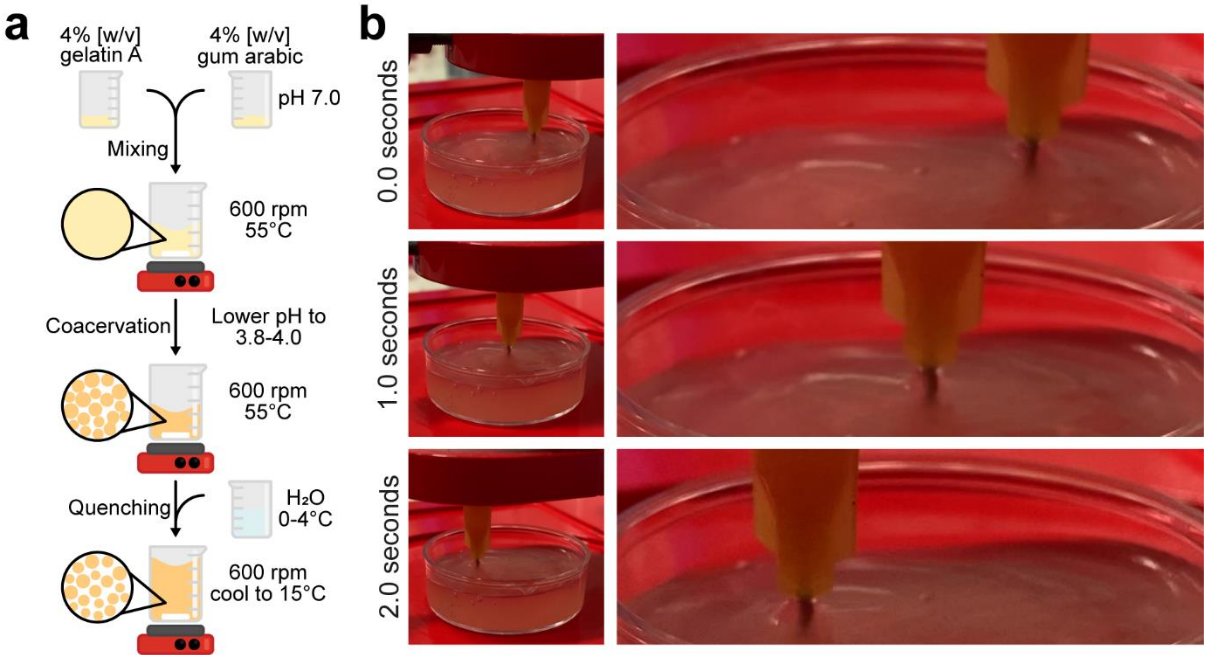


**Supplementary Fig. 1: Compacted microparticles display self-healing properties.**
**a**, Schematic illustration of microparticle generation. Solutions of gelatin type A and gum arabic (both 4% [w/v]) are mixed at 55°C under constant stirring at 600 rpm. Coacervation is induced by lowering the pH of the solution to 3.8–4.0. Microparticles are quenched under constant stirring by the addition of cold water (0–4°C) and further cooling in an ice bath to 15°C. **b**, Still images taken from a video of a printing needle passing through compacted microparticles showing self-healing properties.

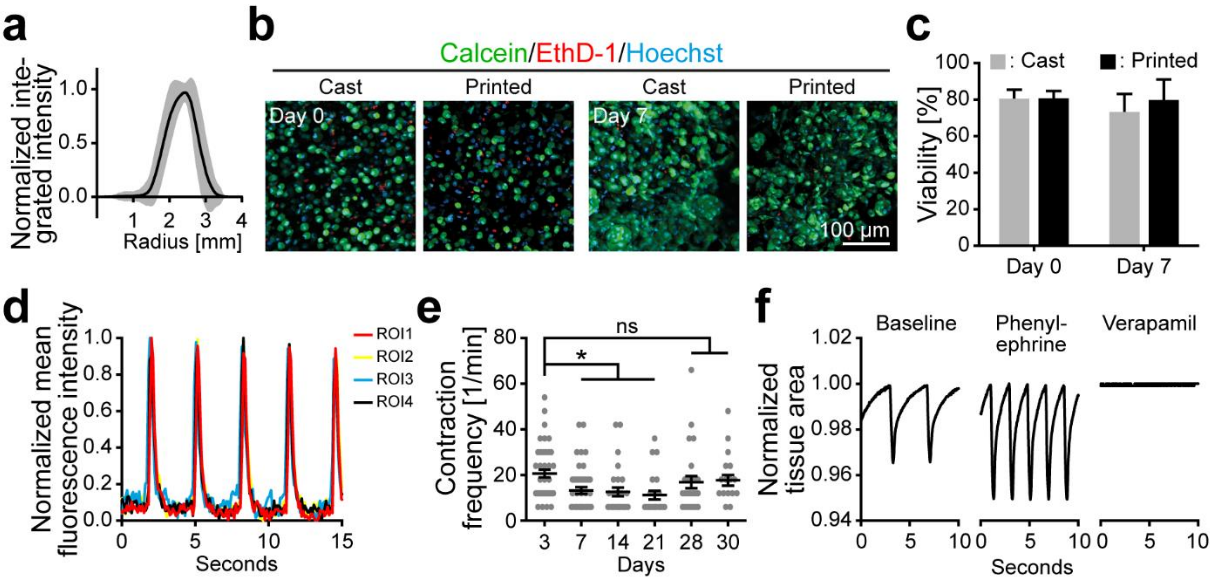


**Supplementary Fig. 2: Printed hiPSC-derived cardiomyocytes form functional cardiac tissues.**

**a**, Radial intensity profile (data is mean ± SD) of rings printed using *CollHA3-3* containing 25 Mio/ml hiPSC-CMs (n = 119). **b**, Representative maximum intensity projections of confocal z-stacks of printed and cast tissues (day 0 and day 7) stained with calcein, ethidium homodimer-1 (EthD-1), and Hoechst to label viable (calcein) and dead (EthD-1) cells. **c**, Quantitative analysis of (**b**). **d** Quantification of calcium flux in printed tissues (day 30) based on normalized mean Fluo-4 fluorescence intensity over multiple contractions in regions of interest in Fig. 3f. **e**, Contraction frequency of printed rings over the course of cultivation from day 3 to 30 (n = 20-45; *: $p < 0.05$). **f**, Representative plots of normalized tissue area over the course of contractions of a printed ring at baseline and subsequently under stimulation with phenylephrine (50 µM) followed by verapamil (1 µM).

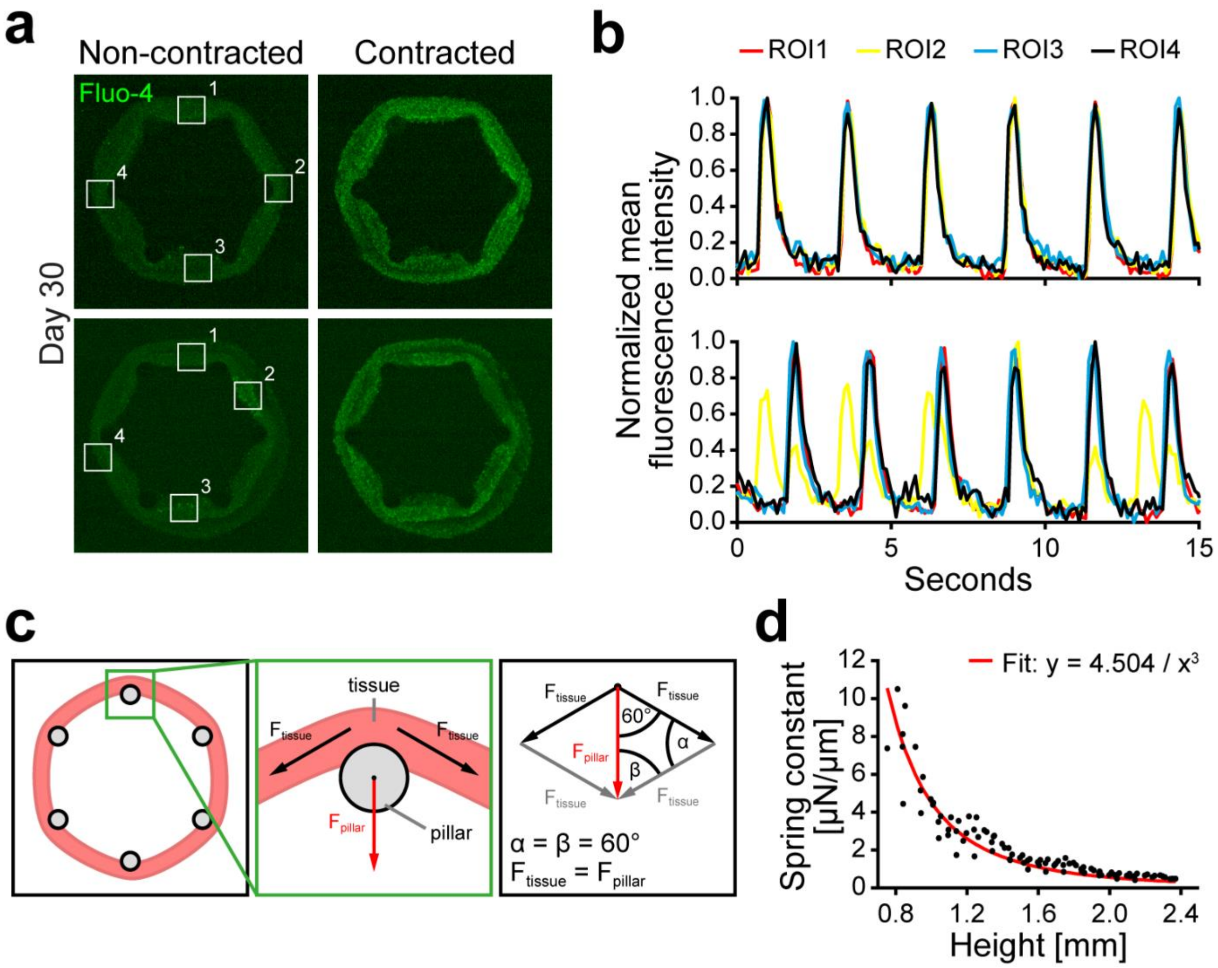


**Supplementary Fig. 3: Printed cardiac tissues exert concentric forces against passive resistance.**

**a**,**b**, Representative examples of calcium flux in non-contracted and contracted printed rings on flexible pillars (day 30) visualized with Fluo-4 (**a**) and its quantification (**b**) based on normalized mean fluorescence intensity over multiple contractions in regions of interest (ROI1–4, white boxes) in (**a**) revealing high tissue synchronicity (upper panel) or the presence of small clusters of cells showing additional “out of rhythm” beats (lower panels, ROI2). **c**, Schematic illustrating the force vectors for printed rings cultured on flexible pillars and resultant force parallelogram used for force calculations. **d**, Calibration curve for determining the spring constants of the pillars. Black dots represent individual measurements of the pillar spring constant $k$ (from the slope of the force-bending relationship) at a given height $h$ above the base. The red line represents the fit of the Euler-Bernoulli beam equation $k = k_0 \, (h/h_0)^{-3}$ to the measurements, with spring constant $k_0$ = 4.5 N/m at a nominal height of $h_0$ = 1 mm above the base. The muscle tissue force $F$ can then be calculated from the pillar bending deflection $d$ according to $F = d \times k$.

**Supplementary Movie Captions**

| | |
|---|---|
| Supplementary movie 1: | beating of a printed ring at day 3 |
| Supplementary movie 2: | calcium handling (Fluo-4) in a printed ring at day 30 |
| Supplementary movie 3: | beating of a printed ring at day 30<br>before and after phenylephrine and verapamil treatment |
| Supplementary movie 4: | beating of a printed ring on a 6 pillar array at day 30 |
| Supplementary movie 5: | calcium handling (Fluo-4) in a printed ring at day 30<br>with high synchronicity |
| Supplementary movie 6: | calcium handling (Fluo-4) in a printed ring at day 30<br>with small areas of asynchronicity |
| Supplementary movie 7: | beating of a printed ring on a 6-pillar array at day 30<br>before and after phenylephrine and verapamil treatment |
| Supplementary movie 8: | beating of a printed ventricle at day 42 |
| Supplementary movie 9: | beating of a printed ventricle at day 100 |
| Supplementary movie 10: | beating of a printed ventricle at day 30<br>before and after phenylephrine stimulation |
